# Single-sequence protein structure prediction using language models from deep learning

**DOI:** 10.1101/2021.08.02.454840

**Authors:** Ratul Chowdhury, Nazim Bouatta, Surojit Biswas, Charlotte Rochereau, George M. Church, Peter K. Sorger, Mohammed AlQuraishi

## Abstract

AlphaFold2 and related systems use deep learning to predict protein structure from co-evolutionary relationships encoded in multiple sequence alignments (MSAs). Despite dramatic, recent increases in accuracy, three challenges remain: (i) prediction of orphan and rapidly evolving proteins for which an MSA cannot be generated, (ii) rapid exploration of designed structures, and (iii) understanding the rules governing spontaneous polypeptide folding in solution. Here we report development of an end-to-end differentiable recurrent geometric network (RGN) able to predict protein structure from single protein sequences without use of MSAs. This deep learning system has two novel elements: a protein language model (AminoBERT) that uses a Transformer to learn latent structural information from millions of unaligned proteins and a geometric module that compactly represents C_α_ backbone geometry. RGN2 outperforms AlphaFold2 and RoseTTAFold (as well as trRosetta) on orphan proteins and is competitive with designed sequences, while achieving up to a 10^6^-fold reduction in compute time. These findings demonstrate the practical and theoretical strengths of protein language models relative to MSAs in structure prediction.

## INTRODUCTION

Predicting 3D protein structure from amino acid sequence is a grand challenge in biophysics. Progress has long relied on physics-based methods that estimate energy landscapes and dynamically fold proteins within these landscapes^1–4^. A decade ago, the focus shifted to extracting residue-residue contacts from co-evolutionary relationships embedded in multiple sequence alignments (MSAs)^5^ **(Supplementary Figure 1)**. Algorithms such as the first AlphaFold^6^ and trRosetta^7^ use deep neural networks to generate distograms that guide classic physics-based folding engines. These algorithms perform substantially better than algorithms based on physical energy models alone. More recently, the outstanding performance of AlphaFold2^8^ on a wide range of protein targets in the recent CASP14 prediction challenge shows that when MSAs are available, machine learning (ML)-based methods can predict protein structure with sufficient accuracy to complement X-ray crystallography, cryoEM, and NMR as a practical means to determine structures of interest.

**Figure 1.**
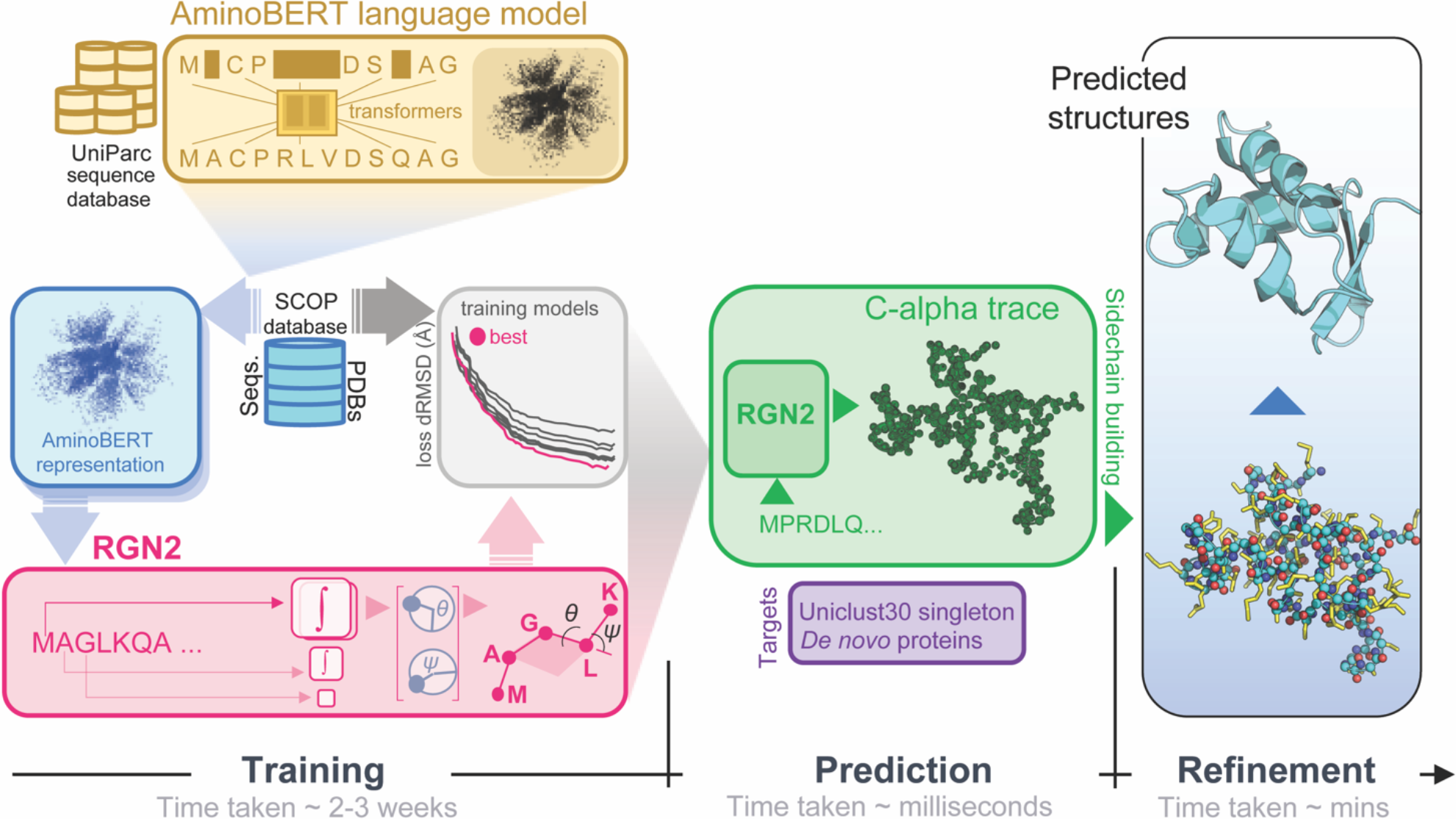
Organization and application of RGN2. RGN2 combines a Transformer-based protein language model (AminoBERT) with a recurrent geometric network that utilizes Frenet-Serret frames to generate the backbone structure of a protein. Placement of side chain atoms and refinement of hydrogen-bonded networks are subsequently performed using the Rosetta energy function.

Predicting the structures of single sequences using ML nonetheless remains a challenge: the requirement in AlphaFold2 for co-evolutionary information from MSAs makes it less performative with proteins that lack sequence homologs, currently estimated at ~20% of all metagenomic protein sequences^9^ and ~11% of eukaryotic and viral proteins^10^. Applications such as protein design and quantifying the effects of sequence variation on function^11^ or immunogenicity^12^ also require single-sequence structure prediction. More fundamentally, the physical process of a polypeptide folding in solution is driven solely by the chemical properties of that chain and its interaction with solvent (excluding, for the moment, proteins that require folding co-factors). An algorithm that predicts sequence directly from a single sequence is—like energy-based folding engines^1–4^—closer to the real-word process than an algorithm that uses MSAs. Thus, we speculate that ML-based single sequence prediction may ultimately provide new understanding of protein biophysics.

Structure prediction algorithms that are fast and low-cost are of great practical value because they make efficient exploration of sequence space possible, particularly in design applications^13^. State of the art MSA-based predictions for large numbers of long proteins can incur substantial costs when performed on the cloud. Reducing this cost would enable many practical applications in enzymology and therapeutics including designing new functions^14–16^, raising thermostability^17^, altering pH sensitivity^18^, and increasing compatibility with organic solvents^19^. Efficient and accurate structure prediction is also valuable in the case of orphan proteins, many of which are thought to play a role in lineage-specific adaptations and are taxonomically restricted. OSP24, for example, is an orphan virulence factor for the wheat pathogen *F. graminearum* that controls host immunity by regulating proteasomal degradation of a conserved signal transduction kinase^20^. It is one of many orphan genes found in fungi, plants insects and other organisms^21^ for which no MSA is available.

We have previously described an end-to-end differentiable, ML-based recurrent geometric network (hereafter RGN1)^22^ that predicts protein structure from position-specific scoring matrices (PSSMs) derived from MSAs; related end-to-end approaches have since been reported^23–25^. RGN1 did not rely on the co-evolutionary information present in MSAs but a requirement for PSSMs necessitates that multiple homologous sequences be available. Here, we describe a new end-to-end differentiable system, RGN2 (**Figure 1**), that predicts protein structure from single protein sequences by using a protein language model (AminoBERT). Such models were first developed for natural language processing (NLP) as a means to extract semantic information from a sequence of words^26^. In the context of proteins, AminoBERT aims to capture the latent information in a string of amino acids that implicitly specifies protein structure. RGN2 also makes use of a natural way of describing polypeptide geometry that is inherently rotationally- and translationally-invariant at the level of the polypeptide as a whole. This involves using the Frenet-Serret formulas to embed a reference frame at each C_α_ carbon; the backbone is then easily constructed by a series of transformations. In this paper we describe the implementation and training of AminoBERT, the use of Frenet-Serret formulae in RGN2, and a performance assessment for natural and designed proteins with no significant sequence homologs. We find that RGN2 consistently outperforms AlphaFold2 (AF2)^8^ and RoseTTAFold (RF)^27^ on naturally occurring orphan proteins without known homologs, and is competitive on *de novo* designed proteins (RGN2 outperforms trRosetta in both cases). While RGN2 is not as performant as MSA-based methods for proteins that permit MSAs, it is up to six orders of magnitude faster, enabling efficient exploration of sequence and structure landscapes.

## RESULTS

### RGN2 and AminoBERT models

RGN1^22^ processed a protein PSSM through a recurrent neural network that implicitly learned PSSM-structure relationships. These relationships were parameterized as torsion angles between adjacent residues making it possible to sequentially position the protein backbone in 3D space (the backbone is described by *N*, *C*_*α*_, and *C′* atoms). All RGN1 components were differentiable and the system could therefore be optimized from end to end to minimize prediction error (as measured by distance-based root mean squared deviation; dRMSD). The RGN2 model described here involves two primary innovations. First, it uses amino acid sequence itself as the primary input as opposed to a PSSM, making it possible to predict structure from a single sequence. In the absence of a PSSM or MSA, latent information on the relationship between protein sequence (as a whole) and 3D structure is captured using a protein language model we term AminoBERT. Second, rather than describe the geometry of protein backbones as a sequence of torsion angles, RGN2 uses a simpler and more powerful approach based on the Frenet-Serret formulas; these formulas describe motion along a curve using the reference frame of the curve itself. This approach to protein geometry is inherently translationally- and rotationally-invariant. We optionally refine predicted structures using a Rosetta-based protocol^28^ that imputes the backbone and side-chain atoms. Refinement is first performed in torsion space to optimize side-chain conformations and eliminate clashes and then in Cartesian space using quasi-Newton-based energy minimization. This protocol is non-differentiable but improves the quality of predicted structures.

Language models were originally developed for natural language processing and operate on a simple but powerful principle: they acquire linguistic understanding by learning to fill in missing words in a sentence, akin to a sentence completion task in standardized tests. By performing this task across large text corpora, language models develop powerful reasoning capabilities. The Bidirectional Encoder Representations from Transformers (BERT) model^29^ instantiated this principle using Transformers, a class of neural networks in which attention is the primary component of the learning system^30^. In a Transformer, each token in the input sentence can “attend” to all other tokens through the exchange of activation patterns corresponding to the intermediate outputs of neurons in the neural network. In AminoBERT we utilize the same approach, substituting protein sequences for sentences and using amino acid residues as tokens. We train a 12-layer Transformer using ~250 million natural protein sequences obtained from the UniParc sequence database^31^. To enhance the capture of information in full protein sequences we introduce two training objectives not part of BERT or previously reported protein language models^26,32–36^. First, 2-8 contiguous residues are masked simultaneously in each sequence making the reconstruction task harder and emphasizing learning from global rather than local context. Second, *chunk permutation* is used to swap contiguous protein segments; chunk permutations preserve local sequence information but disrupt global coherence. Training AminoBERT to identify these permutations is another way of encouraging the Transformer to discover information from the protein sequence as whole. The AminoBERT module of RGN2 is trained independently of the geometry module in a self-supervised manner without fine-tuning (see Methods for details).

In RGN2 we parameterize backbone geometry using the discrete version of the Frenet-Serret formulas for one-dimensional curves^37^. In this parameterization, each residue is represented by its *C*_*α*_ atom and an oriented reference frame centered on that atom. Local residue geometry is described by a single rotation matrix relating the preceding frame to the current one, which is the geometrical object that RGN2 predicts at each residue position. This rotationally- and translationally-invariant parameterization has two advantages over our use of torsion angles in RGN1. First, it ensures that specifying a single biophysical parameter, namely the sequential *C*_*α*_ − *C*_*α*_ distance of ~3.8Å (which corresponds to a trans conformation) results in only physically-realizable local geometries. This overcomes a limitation of RGN1 which yielded chemically unrealistic values for some torsion angles. Second, it reduces by ~10-fold the computational cost of chain extension calculations, which often dominate RGN training and inference times (see Methods).

RGN2 training was performed using both the ProteinNet12 dataset^38^ and a smaller dataset comprised solely of single protein domains derived from the ASTRAL SCOPe dataset (v1.75)^39^. Since we observed no detectable difference between the two, all results in this paper derive from the smaller dataset as it required less computing time to train.

### Predicting structures of proteins with no homologs

To assess how well RGN2 predicts the structures of orphan proteins having no known sequence homologs, we compared it to AlphaFold2 (AF2)^8^, RoseTTAFold (RF)^27^, and trRosetta^7^, currently the best publicly available methods. As a test set, we used 222 orphan proteins that constitute a complete cluster in the Uniclust30 dataset^40^ (*i.e.,* they have no homologs). Of these, 196 have structures available in the Protein Databank (PDB)^41^ but are not part of the training sets used for RGN2 or trRosetta (Methods). However, more than 95% of these sequences are included in the training sets for AF2 and RF, which may result in an overestimate of their accuracy. We predicted the structures of these orphan proteins using all methods and assessed accuracy with respect to experimentally-determined structures (**Figure 2A**) using dRMSD and GDT_TS (the global distance test, which roughly captures the fraction of the structure that is correctly predicted). We found that RGN2 outperformed AF2, RF, and trRosetta on both metrics in 51%, 56%, and 63% of cases, respectively (these correspond to the top-left quadrant in **Figure 2B** and **Supplementary Figure 2**). In 10%, 21%, and 4% of cases, AF2, RF, and trRosetta outperformed RGN2 on both metrics, respectively; split results were obtained in the remaining cases. When we computed differences in error metrics among methods, we found that RGN2 outperformed AF2 (and RF; trRosetta) by an average ΔdRMSD of 1.85Å (3.26Å; 4.36Å) and ΔGDT_TS of 3.51 (6.87; 10.21).

**Figure 2.**
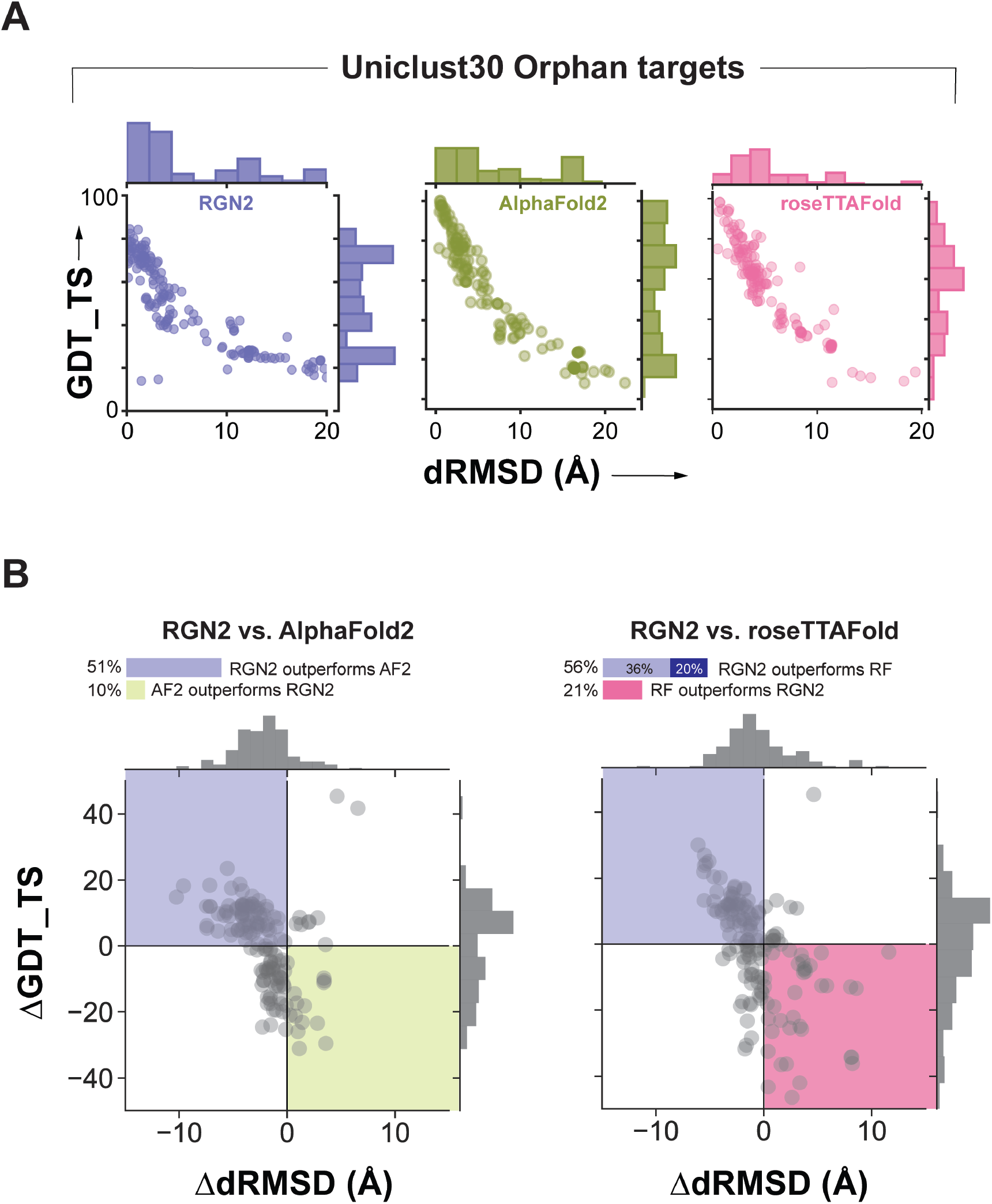
**(A)** Absolute performance metrics for RGN2 (purple), AF2 (green), and RF (pink) across 196 orphan proteins lacking known homologs. Outliers in RGN2 plot correspond to short targets where a few poorly predicted residues substantially lower GDT_TS. **(B)** Differences in prediction accuracy between RGN2 and AF2 / RF are shown for the 196 orphan proteins, using dRMSD and GDT_TS as metrics. RF failed to converge for 40 targets, yielding no predictions. Points in top-left quadrant correspond to targets with negative ΔdRMSD and positive ΔGDT_TS, *i.e.,* where RGN2 outperforms the competing method on both metrics, and vice-versa for the bottom-right quadrant. The other two quadrant (white) indicate targets where there is no clear winner as the two metrics disagree.

To investigate the basis for these differences in performance we determined the fraction of each secondary structure element (alpha-helix, beta-strand, hydrogen-bonded turn, 3/10-helix, bend, beta-bridge, and 5-helix) by applying the DSSP algorithm^42^ to the PDB structures in the orphan protein test set (**Figure 3A** and **Supplementary Figures 2**, **3**, and **4)**. We found that RGN2 outperformed all other methods on proteins rich in bends and on hydrogen-bonded turns interspersing helices, while other methods—AF2 in particular—better predicted targets with high fractions of beta-strand and beta-bridges (such as hairpins). Performance on the remaining ~30% targets was split between RGN2 and competing methods (**Figure 2B**). We also examined performance as a function of protein length and observed that RGN2 generally outperformed AF2 on longer proteins. One possible reason for these findings is that alpha helix formation is strongly driven by local sequence preferences, making helices more readily detectable from single sequences by RGN2.

**Figure 3.**
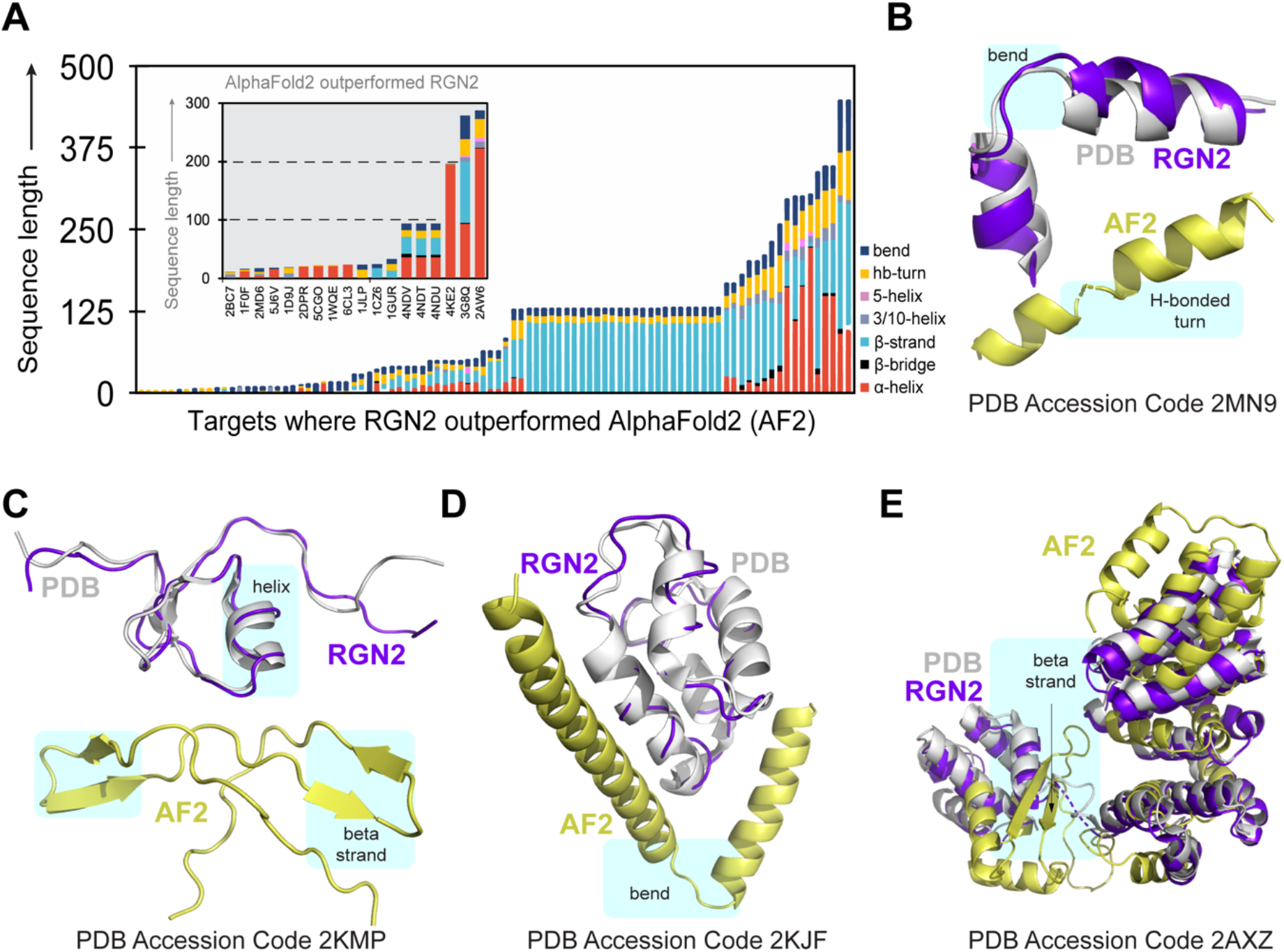
**(A)** Stacked bar chart shows the relative fractions of secondary structure elements in orphan proteins where RGN2 outperforms AF2 (the 18 proteins on which AF2 outperforms RGN2 are shown in the gray inset). Bar height indicates protein length. **(B-E)** Alpha helical targets with bends or turns between helical domains tend to be better predicted by RGN2. AF2, as shown in 2KMP and 2AXZ, often predicts spurious beta strands for orphan proteins.

In **Figure 3B-E** we show examples of structures for which RGN2 outperformed AF2. PDB structure 2MN9 (76% alpha helical) has two short alpha helices (**Figure 3B**) connected by a hydrogen-bonded turn and a polypeptide bend. RGN2 correctly predicts the challenging, less-structured bends and turns in this protein, yielding a 37-point gain in GDT_TS (ΔdRMSD > 6Å) over AF2. Similar trends were observed for structures 2KMP, 2KJF, and 2AXZ (**Figure 3C-E**). 2AXZ is another protein with a single stranded helical bundle connected by bends, albeit one that is longer than 2KJF (305 residues as opposed to 60 residues). AF2 predicted the majority of helical domains in 2AXZ accurately, but a 24 amino acid fragment from Y17 through R40 was incorrectly predicted to be a beta-beta hairpin; in contrast, RGN2 correctly predicted this polypeptide sequence to comprise two helices connected by a bend. This contributed to a ~17-point gain in GDT_TS and 5.3Å increase in dRMSD by RGN2 relative to AF2.

### Predicting the structures of de novo (designed) proteins

To determine RGN2 accuracy on designed proteins we used a set of 35 proteins generated *de novo* using the Rosetta energy function; such proteins are expected to be well-suited to prediction by RF and trRosetta. The designed proteins primarily have therapeutic applications ranging from anti-microbial activity (1D9J and 6CL3) to blocking of human potassium channels (1WQD). As before, we assessed prediction accuracy using dRMSD and GDT_TS. We found that RGN2 outperformed AF2, RF, and trRosetta on both metrics in 17%, 26%, and 45% of cases, respectively (**Supplementary Figure 5**) but underperformed on both metrics in 49%, 54%, 7% of cases (**Figure 4**). On average, AF2 exhibited the best overall performance; however, relative to RF, RGN2 exhibited a 3.5Å better average dRMSD and marginally worse (lower) average GDT_TS of 3.2 (**Figure 4B**). When we grouped proteins by secondary structure content (**Figure 5**) we found that AF2 outperformed RGN2 for proteins such as 6D0T, 87% of which is a beta sheet arranged in a radially symmetric manner. In contrast, RGN2 outperformed AF2 (and RF) on structures such as 6TJ1 that comprise alpha helices held in place by hydrogen-bonded turns. We conclude that RGN2 can learn sequence-structure relationships for *de novo* regions of protein space, but that beta sheet prediction from single sequences remains a challenge. The superior performance of RGN2 relative to AF2 on a non-trivial fraction of *de novo* proteins also suggests that the two methods may be complementary and a hybrid model may outperform either one alone.

**Figure 4.**
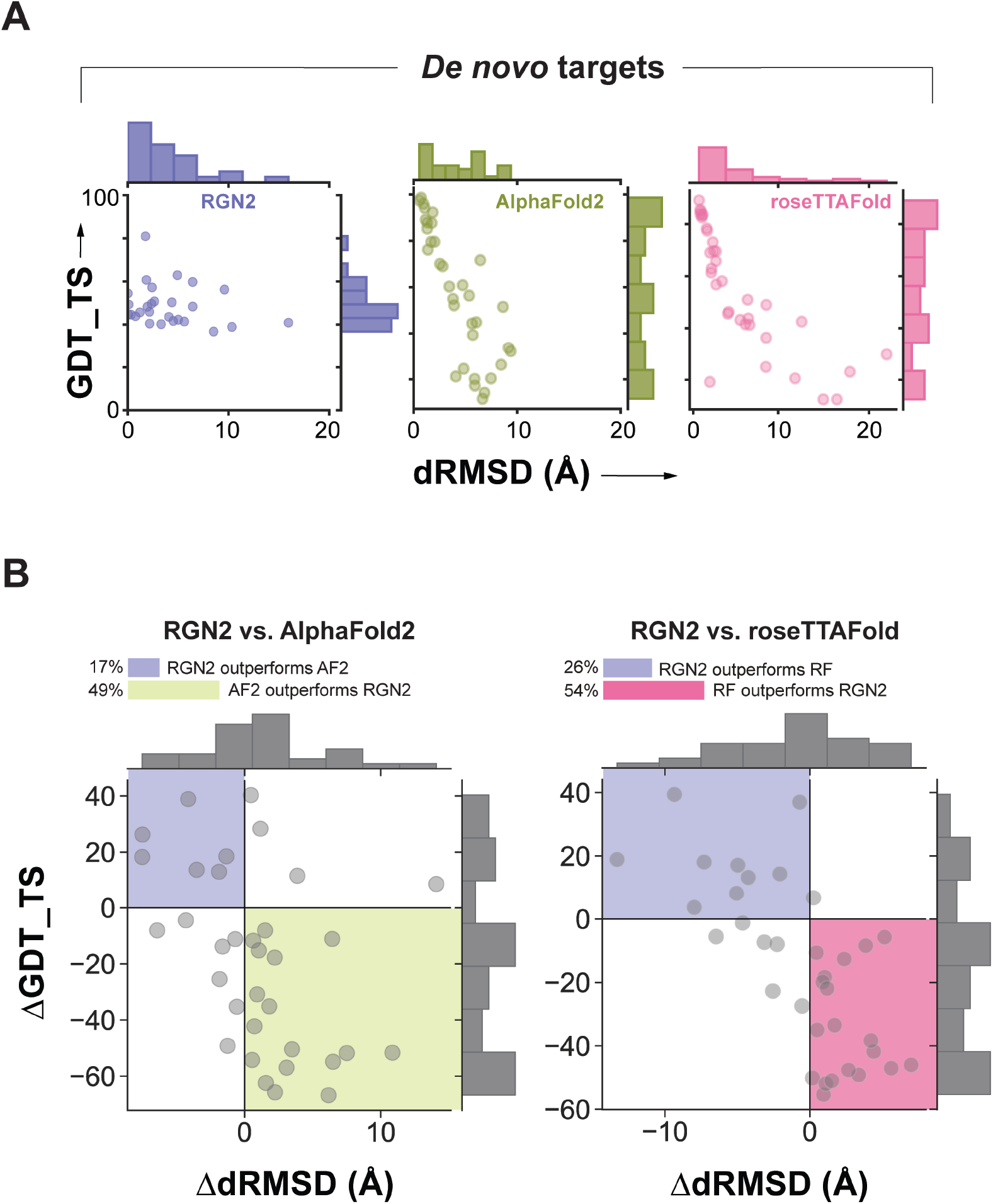
**(A)** Absolute performance metrics for RGN2 (purple), AF2 (green), and RF (pink) across 35 *de novo* designed proteins with no known homologs. **(B)** Differences in prediction accuracy between RGN2 and AF2/RF are shown for these 35 proteins, using dRMSD and GDT_TS as metrics. Points in top-left quadrant correspond to targets with negative ΔdRMSD and positive ΔGDT_TS, *i.e.,* where RGN2 outperforms the competing method on both metrics, and vice-versa for the bottom-right quadrant. The other two quadrant (white) indicate targets where there is no clear winner as the two metrics disagree.

**Figure 5.**
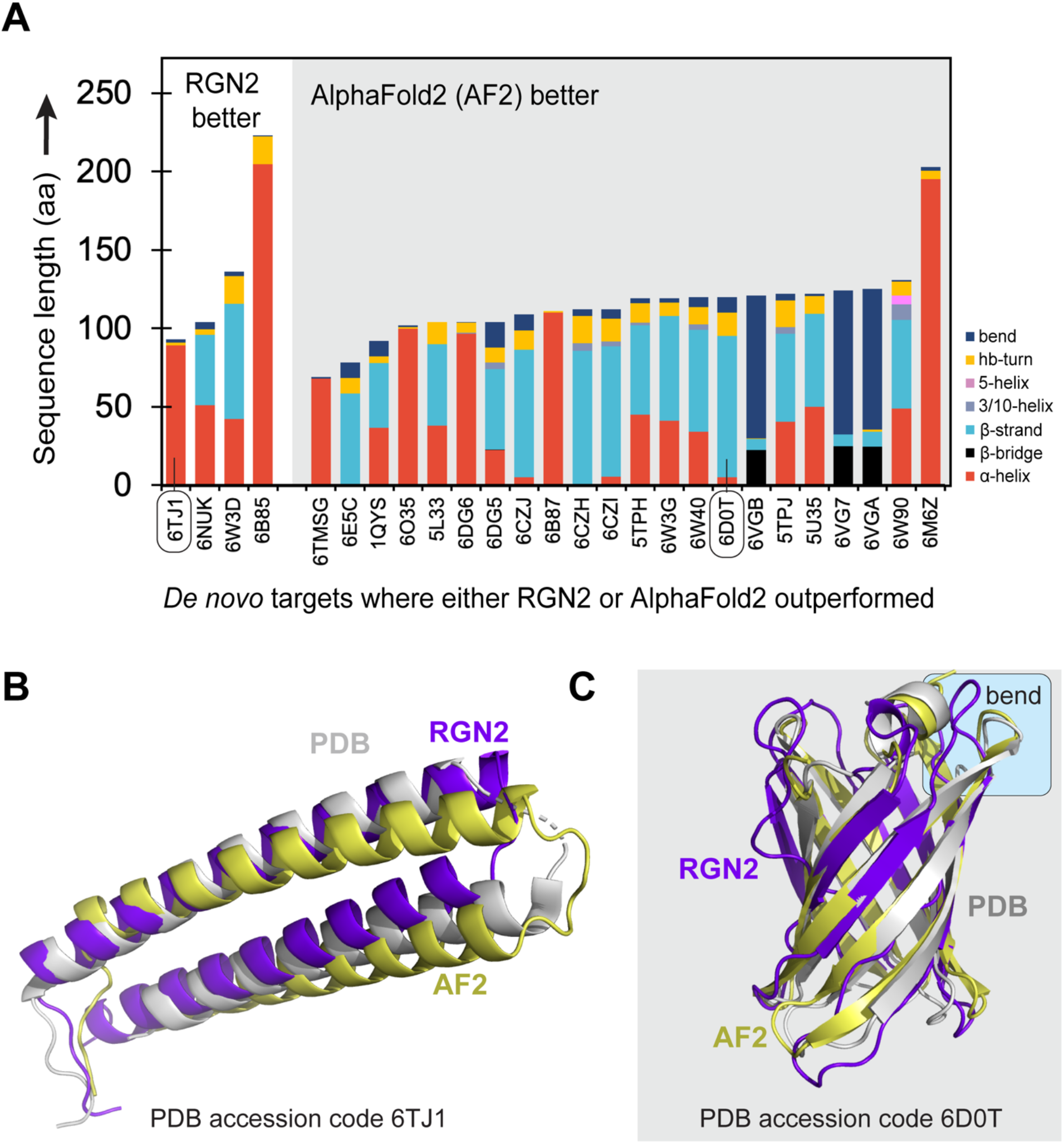
**(A)** Stacked bar chart shows the relative fractions of secondary structure elements in 35 *de novo* proteins. Bar height indicates protein length. AF2 outperforms RGN2 on proteins with high beta-sheet content. **(B)** Alpha helical target 6TJ1 does not have a bend, but AF2 predicts a spurious bend while the RGN2 prediction is closer to the experimental structure. **(C)** 6D0T, a combination of an hour-glass shaped beta barrel connected by disordered loops, is better predicted by AF2.

### RGN2 prediction speed

Rapid prediction of protein structure is essential for tasks such as protein design and analysis of allelic variation or disease mutations. By virtue of being end-to-end differentiable, RGN2 predicts unrefined structures using fast neural network operations and does not require physics-based conformational sampling to assemble a folded chain. Moreover, by virtue of operating directly on single sequences, RGN2 avoids expensive MSA calculations. To quantify these benefits, we compared the speed of RGN2 and other methods on orphan and *de novo* proteins datasets of varying lengths (breaking down computation time by prediction stage; **Table 1**). In MSA-based methods, MSA generation scaled linearly with MSA depth (*i.e.,* the number of homologous sequences used) whereas distogram prediction (by trRosetta) scaled quadratically with protein length. AF2 predictions scale cubically with protein length. In contrast, RGN2 scales linearly with protein length and both template-free and MSA-free implementations of AF2 and RF were >10^5^ fold slower than RGN2. In the absence of post-prediction refinement, RGN2 is up to 10^6^ folds faster, even for relatively short proteins Moreover, predictions for 40 orphan targets (*e.g.,* 2MN9 and 2MOA) failed to converge using MSA or template-free versions of RF, despite being less than 65 residues long on average. Adding physics-based refinement increases compute cost for all methods, but even so RGN2 remains the fastest available method. Of interest, even when MSA generation is discounted, neural network-based inference for AF2 and RF remains much slower than RGN2, inclusive of post-prediction refinement. This gap will only widen for design tasks involving longer proteins, as is increasingly becoming possible, and we interpret it to be a measure of the value of using a protein language model such as AminoBERT in a prediction algorithm.

**Table 1.** Comparison of prediction times between RGN2 and AF2, RF, and trRosetta across 231 targets spanning our orphan and *de novo* protein datasets. RGN2 predictions were performed in batches with maximum permissible batch size set to 128 targets. The trRosetta MSA generation step was not used since none of the targets had known homologous proteins.

| Protein length<br>(L) bins<br>(# residues) | Total<br>targets | Mean<br>protein<br>length<br>(#<br>residues) | Mean trRosetta prediction time<br>per structure (s) |  | Mean<br>AF2<br>prediction<br>time<br>(s) | Mean<br>RF<br>prediction<br>time<br>(s) | Mean<br>RGN2<br>prediction<br>time<br>(ms) | Mean RGN2<br>prediction +<br>refinement<br>time (s) |
| --- | --- | --- | --- | --- | --- | --- | --- | --- |
|  |  |  | Distogram | 3D-Structure |  |  |  |  |
| $0 < L \leq 100$ | 124 | 33.35 | 1868 | 1021 | 227.8 | 346.2 | 2.6 | 140.36 |
| $100 < L \leq 200$ | 69 | 132.68 | 2762 | 1845 | 252.2 | 376.3 | 2.5 | 169.87 |
| $200 < L \leq 300$ | 18 | 240.27 | 2940 | 1698 | 221.6 | 389.4 | 3.6 | 178.41 |
| $300 < L \leq 400$ | 10 | 328.8 | 3511 | 2135 | 226.9 | 388.6 | 5.1 | 202.31 |
| $400 > L$ | 10 | 457.3 | 4003 | 2977 | 235.1 | 386.8 | 5.3 | 226.23 |

## DISCUSSION

RGN2 represents one of the first attempts to use ML to predict protein structure from a single sequence. This is computationally efficient and has many advantages in the case of orphan and designed proteins for which generation of multiple sequence alignment is not possible. RGN2 accomplishes this by fusing a protein language model (AminoBERT) with a simple and intuitive approach to describing C_α_ backbone geometry based on the Frenet-Serret formulation. Whereas most recent advances in ML-based structure prediction have relied on MSAs^5^, AminoBERT learns information from proteins as a whole without alignment and was trained to capture global structural properties by using sequences with masked residues and block permutations. We speculate that the latent space of the language model also captures recurrent evolutionary relationships^43^. The use of Frenet-Serret formulas in RGN2 addresses the requirement that proteins exhibit translational and rotational invariance. From a practical standpoint, the speed and accuracy of RGN2 shows that language models are nearly as effective as MSAs at learning structural information from primary sequence while being able to extrapolate beyond known proteins, allowing for effective prediction of orphan and designed proteins.

Transformers and their embodiment of local and distant attention is a key feature of language models such as AminoBERT. Very large Transformer-based models trained on hundreds of millions and potentially billions of protein sequences are increasingly available^26,35,36^ and the scaling previously observed in natural language applications^44^ makes it likely that the performance of RGN2 and similar methods will continue to improve. AlphaFold2 also exploits attention mechanisms based on Transformers to capture the latent information in MSAs. Similarly, the self-supervised MSA Transformer^45^ uses a related attention strategy that attends to both positions and sequences in an MSA, and achieves state-of-the-art contact prediction accuracy. It is possible that language models will replace MSAs in general use but more likely are new architectures that merge the two. Data augmentation by learning from high-confidence structures in the newly reported AlphaFold Database^8^ is expected to further improve performance. Finally, training on experimental data is likely to be invaluable in selected applications such as predicting structural variation within kinases or G protein-coupled receptors.

We consider RGN2 to be a first step in learning a direct sequence-to-structure map without a requirement for explicit evolutionary information. One limitation of RGN2 as currently implemented is that the immediate output of the recurrent geometric network only constrains local dependencies between C_α_ atoms (curvature and torsion angles) resulting in sequential reconstruction of backbone geometry. Allowing the network to reason directly on arbitrary pairwise dependencies throughout the structure, and using a better inductive prior than learning immediate contacts may further improve the quality of model predictions. Furthermore, as currently implemented, refinement in RGN2 is not yet part of an end-to-end implementation; refinement via a 3D rotationally- and translationally-equivariant neural network would be more efficient and likely yield better quality structures.

It has been known since Anfinsen’s refolding experiments that single polypeptide chains contain the information needed to specify fold^46^. The demonstration that a language model can learn information on structure directly from protein sequences and then guide accurate prediction of an unaligned protein suggests that RGN2 behaves in a manner that is more similar to the actual folding process than MSA-based methods. Language models learned by deep neural networks are readily formulated in a maximum entropy framework^47^ and the physical process of protein folding is also entropically driven, potentially suggesting a means to compare the two. A fusion of biophysical and learning-based perspectives may ultimately prove the key to direct sequence-to-structure prediction from single polypeptides at the accuracy of experimental methods.

## METHODS

### AminoBERT summary

AminoBERT is a 12-layer Transformer where each layer is composed of 12 attention heads. It is trained to distill protein sequence semantics from ~260 million natural protein sequences obtained from the UniParc sequence database^31^ (downloaded May 19, 2019).

During training each sequence is fed to AminoBERT according to the following algorithm:

1. With probability 0.3 select sequence for chunk permutation, and with probability 0.7 select sequence for masked language modeling.
2. If sequence was selected for chunk permutation, then: With probability 0.35 chunk permute, else (with probability 0.65) leave the sequence unmodified.
3. Else if the sequence was selected for masked language modeling, then: With probability 0.3 introduce 0.15 × sequence_length masks into the sequence with clumping, else (with probability 0.7) introduce the same number of masks into the sequence randomly across the length of the sequence (standard masked language modeling).

The loss for an individual sequence (seq) is given by:

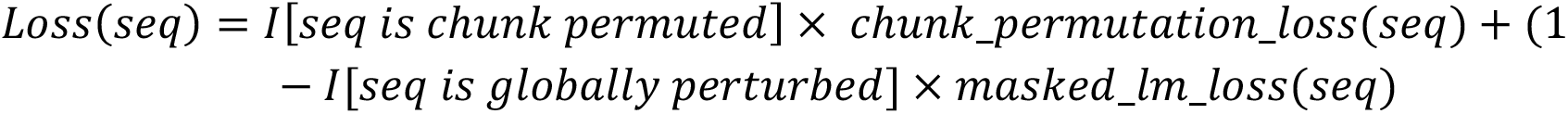

where *I*[x] is the indicator of the event x, and returns 1 if x is true, and 0 if x is false. *Chunk_permutation_loss(seq)* is a standard cross entropy loss reflecting the classification accuracy of predicting whether seq has been chunk permuted. Finally, *masked_lm_loss(seq)* is the standard masked language modeling loss as previously described in Devlin *et al*.^29^. Note, that mask clumping does not affect how the loss is calculated.

Chunk permutation is performed by first sampling an integer *x* uniformly between 2 and 10, inclusively. The sequence is then randomly split into *x* equal-sized fragments, which are subsequently shuffled and rejoined.

Mask clumping is performed as follows:

1. Sample an integer *clump_size* ~ Poisson (2.5) + 1
2. Let *n_mask* = 0.15 × *sequence_length*. Randomly select *n_mask*/*clump_size* positions in the sequence around which to introduce a set of *clump_size* contiguous masks

### AminoBERT architecture

Each multi-headed attention layer in AminoBERT contains 12 attention heads, each with hidden size 768. The output dimension of the feed-forward unit at the end of each attention layer is 3072. As done in BERT^29^, we prepend a [CLS] token at the beginning of each sequence, for which an encoding is maintained through all layers of the AminoBERT Transformer. Each sequence was padded or otherwise clipped to length 1024 (including the [CLS] token).

For chunk permutation classification, the final hidden vector of the [CLS] token is fed through another feed forward layer of output dimension 768, followed by a final feed forward layer of output dimension 2, which are the logits corresponding to whether the sequence is chunk permuted or not. Masked language modeling loss calculations are set up as described in Devlin *et al*. ^29^.

### AminoBERT training procedure

AminoBERT was trained with batch size 3072 for 1,100,000 steps, which is approximately 13 epochs over the 260 million sequence corpus. For our optimizer we used Adam with a learning rate of 1e-4, β1 = 0.9, β2 = 0.999, epsilon=1e-6, L2 weight decay of 0.01, learning rate warmup over the first 20,000 steps, and linear decay of the learning rate. We used a dropout probability of 0.1 on all layers, and used GELU activations as done for BERT. Training was performed on a 512 core TPU pod for approximately one week.

### Geometry module

The geometry of the protein backbone as summarized by the *C*_*α*_ trace can be thought of as a one-dimensional discrete open curve, characterized by a bond and torsion angle at each residue. Following *Niemi et al.*^37^, the starting point for describing such discrete curves is to assign a frame, a triplet of orthonormal vectors, to each *C*_*α*_ atom. If we denote by *r*_*α*_ the vector characterizing the position of a *C*_*α*_ atom at the *i*-th vertex, we could then define a unit tangent vector along an edge connecting two consecutives *C*_*α*_ atoms

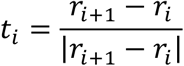

For assigning frames to each *i*-th *C*_*α*_ atom, we need two extra vectors, the binormal and normal vectors defined as follows:

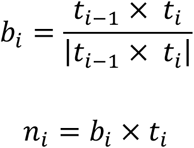

While for a protein (in a given orientation) the tangent vector is uniquely defined, the normal and binormal vectors are arbitrary. Indeed, when assigning frames to each residue we could take any arbitrary orthogonal basis on the normal plane to the tangent vector. Such arbitrariness does not affect our strategy of predicting 3D structures starting from bond and torsion angles.

To derive the equivalent of the Frenet-Serret formulas—which describe the geometry of continuous and differentiable one-dimensional curves—for the discrete case, we need to relate two consecutive frames along the protein backbone in terms of rotation matrices

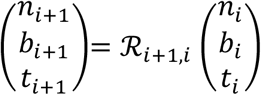

In three-dimensions, rotation matrices are in general parametrized in term of three Euler angles. However, in our case the rotation matrices relating two consecutive frames are fully characterized by only two angles, a bond angle *ψ* and a torsion angle *θ*, as the third Euler angle vanishes, reflecting the following condition *b*_*i*+1_. *t*_*i*_ = 0. We can now write the equivalent of the Frenet-Serret formulas for the discrete case

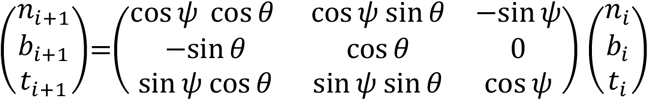

The bond and torsion angles are defined by the following relations

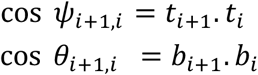

We now turn to backbone reconstruction starting from bond and torsion angles. First, using tangent vectors along the backbone edges, we can reconstruct all *C*_*α*_ atom positions, and thus the full protein backbone in the *C*_*α*_ trace, by using the following relation:

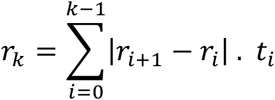

where |*r*_*i*+1_ − *r*_*α*_| is the length of the virtual bonds connecting two consecutive *C*_*α*_ atoms. In most cases, the average virtual bond length is ~ 3.8 Å, which corresponds to trans conformations. In terms of the familiar torsion angles *ϕ, ψ*, and *ω*, those conformations are achieved for *ω* ~ *π*. For cis conformations, mainly involving proline residues, the virtual bond length is ~ 3.0 Å (and it corresponds to *ω* ~ 0). In RGN2, for backbone reconstruction, we impose the condition that the virtual bond length is strictly equal to 3.8 Å, and for reconstructing the backbone we use the following relation:

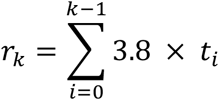

The intuition behind the previous equation is the idea of a moving observer along the protein backbone. We could think of the tangent vector *t*_*i*_ as the velocity of the observer along a given edge, and the constant virtual bond length as the effective time spent for travelling along the edge. The only freedom allowed for such observer is to abruptly change the direction of the velocity vector at each vertex.

The model outputs bond and torsion angles. By centering the first *C*_*α*_ atom of the protein backbone at the origin of our coordinate system, we sequentially reconstruct all the *C*_*α*_ atom coordinates using the following relation:

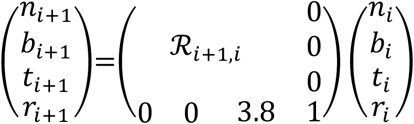

### Data preparation for comparison with trRosetta

Performance of RGN2 was compared against trRosetta across two sets of non-homologous proteins: (i) 196 orphans from the Uniclust30 database^40^, and (ii) 35 *de novo* proteins by Xu *et al*.^48^. Both sets were filtered to ensure no overlap with the training sets of RGN2 and trRosetta. While RGN2 is trained on the ASTRAL SCOPe (v1.75) dataset^39^, trRosetta was trained on a set of 15,051 single chain proteins (released before May 1, 2018).

### Structure prediction with trRosetta, AF2, and RF

Conventional trRosetta-based structure prediction involves first feeding the input sequence through a deep MSA generation step. For orphans and *de novo* proteins without any sequence homologs, the MSA only includes the original query sequence. Next, the MSA is used by the trRosetta neural network to predict a distogram (and orientogram) that captures inter-residue (C_*α*_-C_*α*_ and C_*β*_-C_*β*_) distances and orientations. This information is subsequently utilized by a final Rosetta-based refinement module. This module first threads a naïve sequence of polyalanines of length equaling the target protein that maximally obeys the distance and orientation constraints. After side-chain imputation that reflects the original sequence, multiple steps including clash elimination, rotamer repacking, and energy minimization are performed to identify the lowest energy structure.

AF2 and RF predictions did not require MSAs since our target proteins don’t have homologs and so we made our predictions using their respective official Google Colab notebooks.

### Structure refinement in RGN2

Raw predictions from RGN2 contain a single C*α* trace of the target protein. After performing a local internal coordinate building step to generate the backbone and side-chain atoms corresponding to the target sequence, we use Rosetta-based refinement to finetune the structure. This refinement comprises hybrid optimization of side chains using five invocations of energy minimization in torsional space followed by a single step of quasi-Newton all-atom minimization in Cartesian space (using the *FastRelax* protocol of RosettaScripts^49^). An optional *CartesianSampler*^49^ mover step can be added to further correct local strain density in the predicted model. The six-step *FastRelax* protocol is repeated for 300 cycles for each target. Finally, 100 cycles of coarse-grained, fast minimization using *MinMover*^49^ is applied to obtain the predicted structure.

## Acknowledgements

We gratefully acknowledge the support of the NVIDIA Corporation for the donation of GPUs used for this research. This work is supported by the DARPA PANACEA program grant HR0011-19-2-0022 and NCI grant U54-CA225088 to PKS.

## Author Contributions

R.C., N.B., S.B., and M.A. conceived of and designed the study. R.C. developed the refinement module and performed all analyses. N.B. developed the geometry module and trained RGN2 models. S.B. developed and trained the AminoBERT protein language model and helped integrate its embeddings within RGN2. C.R. trained several RGN2 models and performed RF predictions. P.K.S. and G.M.C. supervised the research and provided funding. N.B., S.B., M.A. and P.K.S wrote the manuscript and all authors discussed the results and edited the final version.

