## Supplementary Figures for "Single-sequence protein structure prediction using language models from deep learning"

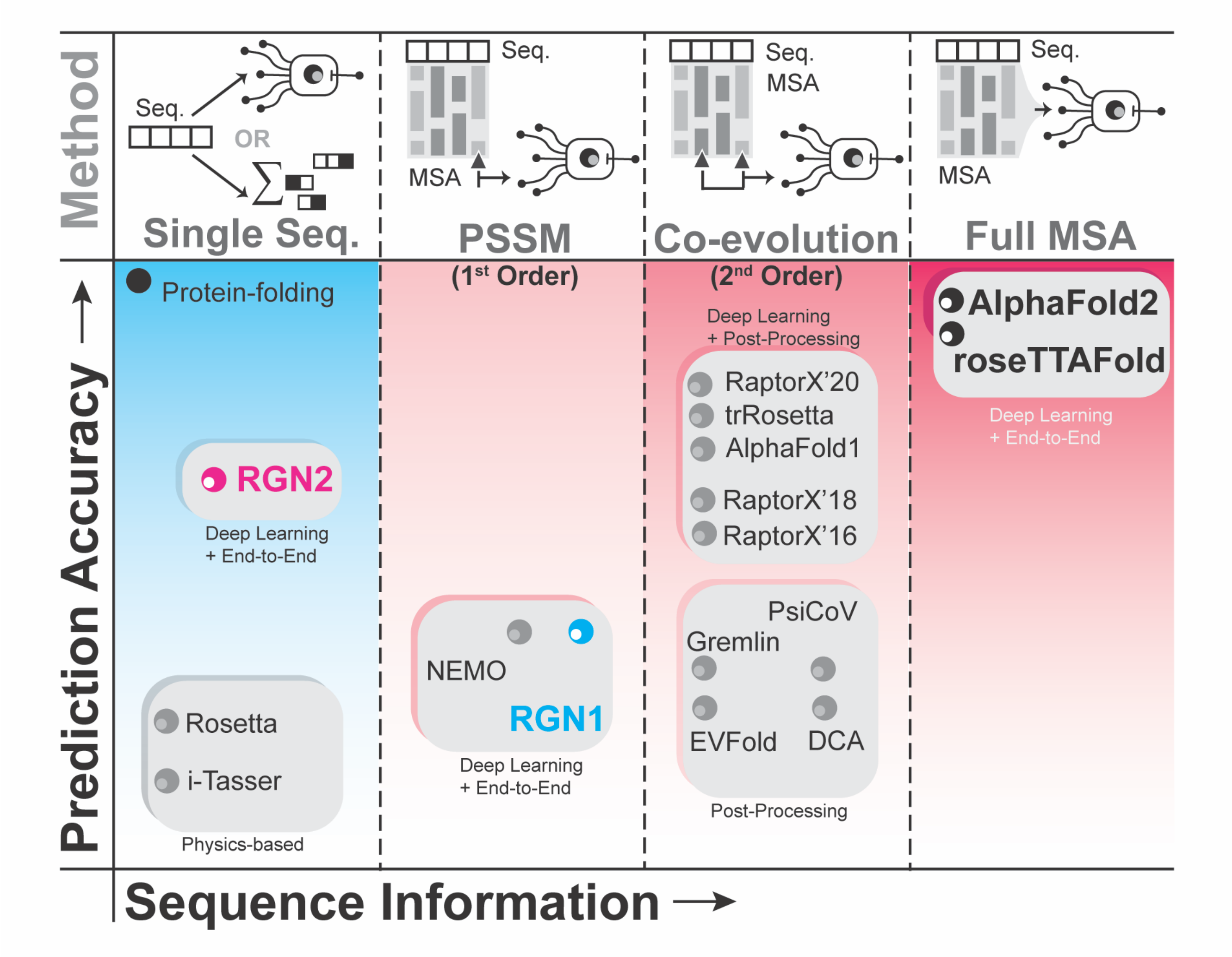


**Supplementary Figure 1. Landscape of protein structure prediction algorithms.** "Physics-based" methods such as Rosetta (lower left) estimate energy landscapes to fold single protein sequences (with or without templates) but they are compute-intensive and not currently high-throughput. Most machine learning-based methods utilize co-evolution information captured by MSAs and PSSMs, which yields a dramatic increase in prediction accuracy. This is exemplified by methods such as RaptorX and AlphaFold2. RGN2 (upper left) revisits the problem of folding single sequences and exhibits superior performance when predicting the structures of orphan or designed proteins for which MSAs cannot be computed.

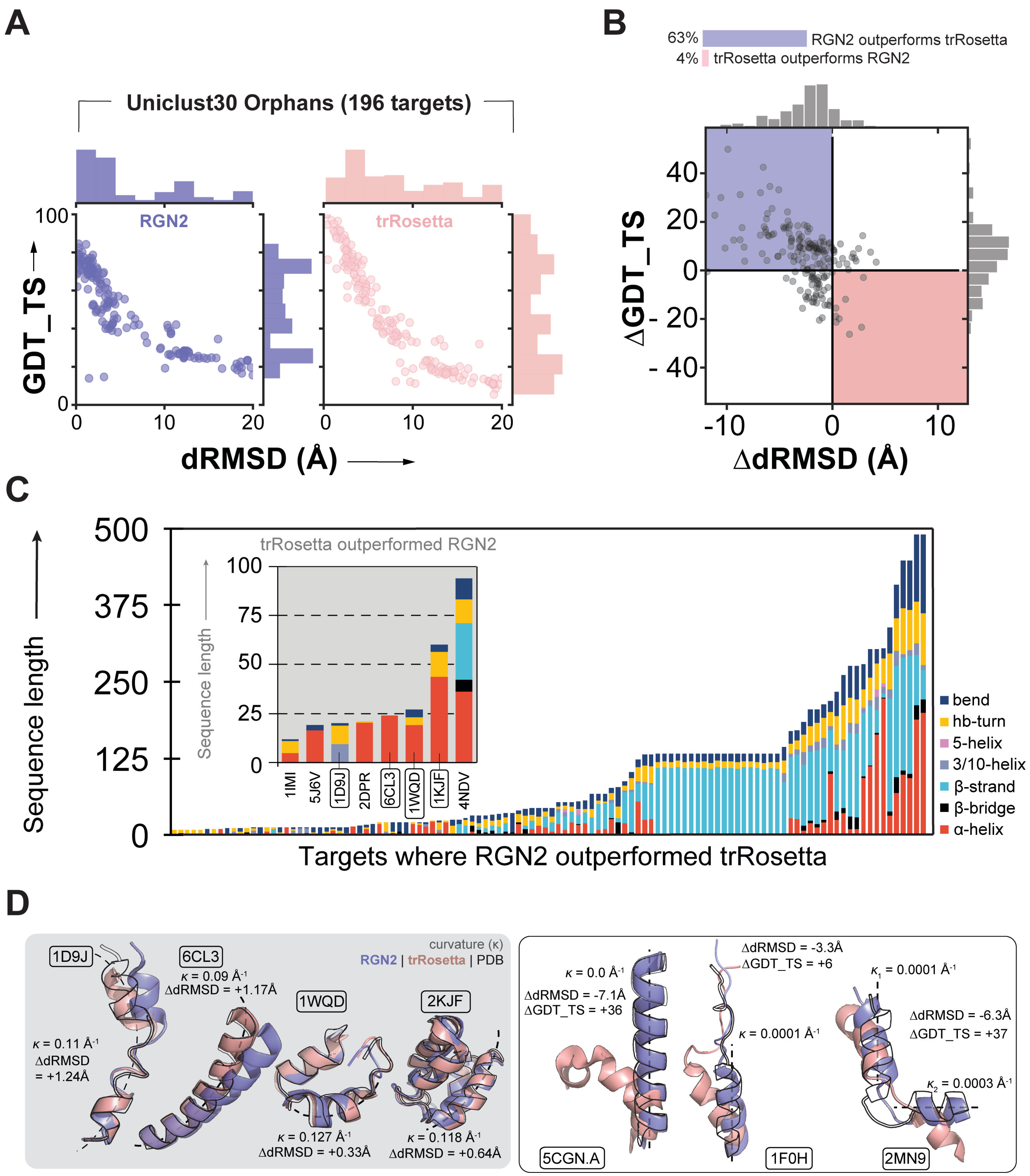


**Supplementary Figure 2. (A)** Absolute performance metrics for RGN2 (purple) and trRosetta (salmon) across two sets of proteins that lack known homologs. Outlier data points for RGN2 plot reflect short targets where erroneous placement of a few residues led to poor predictions. **(B)** Differences between RGN2 and trRosetta dRMSD values were computed for 196 orphan proteins. Points in left-top quadrant correspond to entries with negative ΔdRMSD and positive ΔGDT_TS, *i.e.,* where RGN2 outperforms trRosetta, and vice-versa for the right-bottom quadrant. The other two quadrants (white) indicate targets where there was no clear winner. Only datapoints with <12Å dRMSD are shown—additional entries exist (>12Å dRMSD) but all lie in the quadrant where RGN2 outperforms trRosetta. **(C)** Stacked bar chart shows the relative fractions of different secondary structure elements in orphan proteins where RGN2 outperformed trRosetta (the eight entries where trRosetta outperformed RGN2 are shown in the gray inset). The total height of each bar indicates protein length. **(D)** (Left) Alpha helical targets with bends in the helix, *i.e.,* non-zero curvature (inverse of radius of curvature), tend to be better predicted by trRosetta. (Right) Alpha helical domains lacking curvature, even if linked by unstructured bends and hydrogen-bonded turns, tend to be better predicted by RGN2.

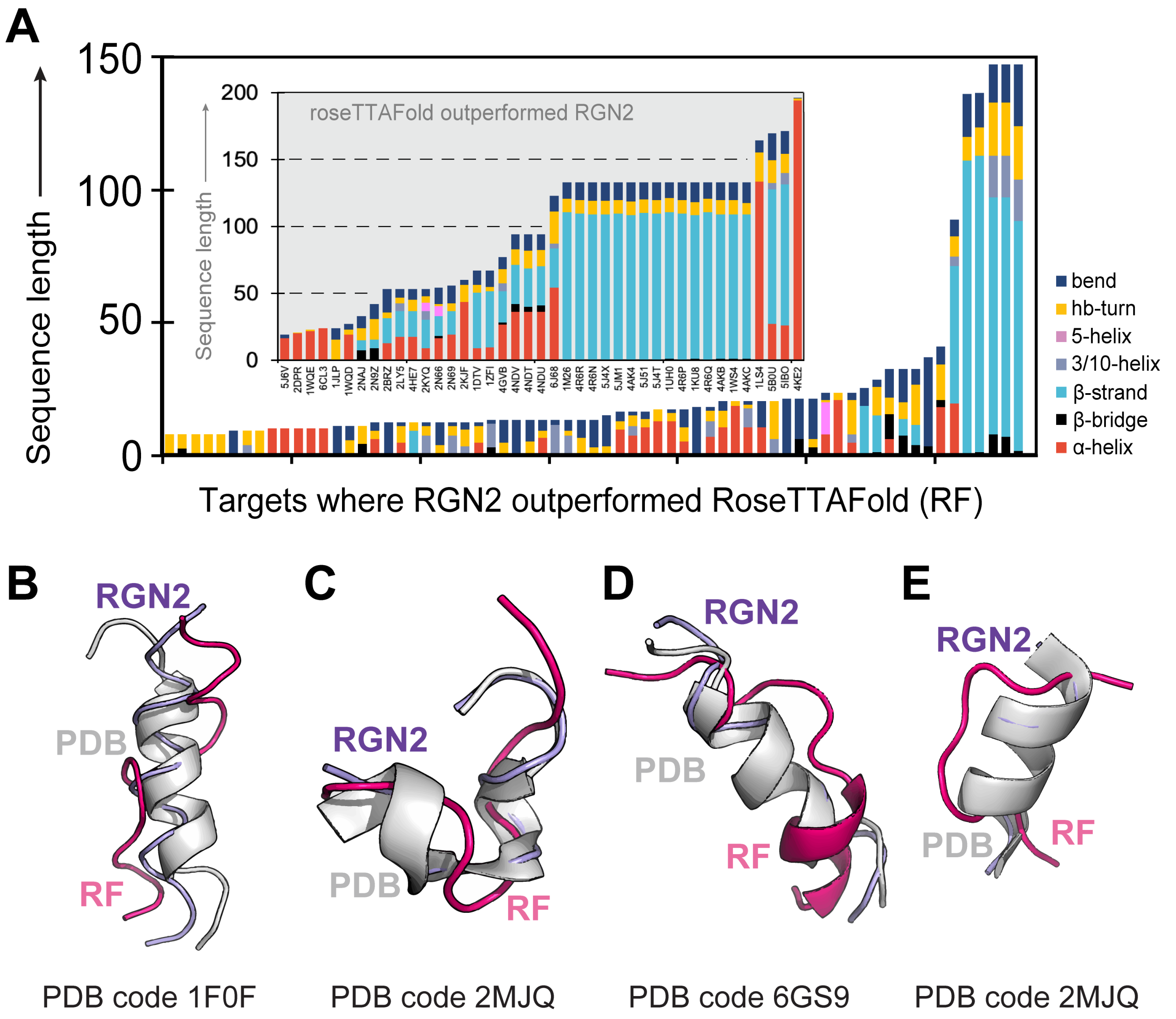


**Supplementary Figure 3. (A)** Stacked bar chart shows the relative fractions of different secondary structure elements in orphan proteins where RGN2 outperformed RF (the proteins where RF outperformed RGN2 are shown in the gray inset.). The total height of each bar indicates protein length. **(B-E)** Secondary structures tend to be better predicted by RGN2. The overall gains in dRMSD and GDT_TS indicate substantially higher prediction accuracy by RGN2 on these proteins.

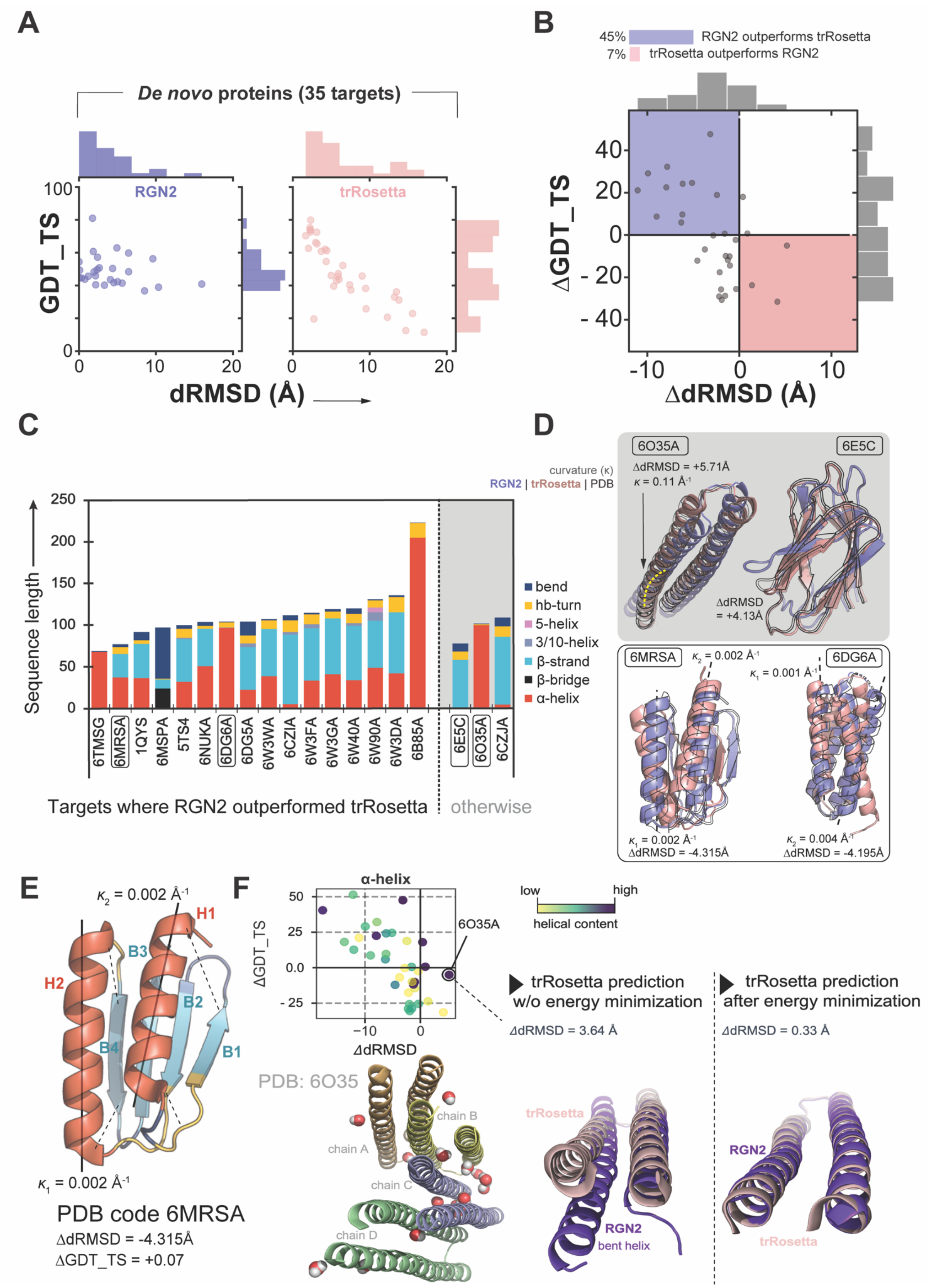

A
De novo proteins (35 targets)
RGN2
trRosetta
GDT_TS
100
0
0
10
20
dRMSD (Å)
B
45%
RGN2 outperforms trRosetta
7%
trRosetta outperforms RGN2
ΔGDT_TS
40
20
0
- 20
- 40
-10
0
10
ΔdRMSD (Å)
C
curvature (κ)
RGN2 | trRosetta | PDB
Sequence length
250
200
150
100
50
0
bend
hb-turn
5-helix
3/10-helix
β-strand
β-bridge
α-helix
6TMSG
6MRSA
1QYS
6MSPA
5TS4
6NUKA
6DG6A
6DG5A
6W3WA
6CZIA
6W3FA
6W3GA
6W40A
6W90A
6W3DA
6B85A
6E5C
6O35A
6CZJA
Targets where RGN2 outperformed trRosetta
otherwise
D
6O35A
6E5C
ΔdRMSD = +5.71Å
κ = 0.11 Å-1
ΔdRMSD
= +4.13Å
6MRSA
6DG6A
κ2 = 0.002 Å-1
κ1 = 0.001 Å-1
κ1 = 0.002 Å-1
ΔdRMSD = -4.315Å
κ2 = 0.004 Å-1
ΔdRMSD = -4.195Å
E
κ2 = 0.002 Å-1
H1
H2
B3
B2
B4
B1
κ1 = 0.002 Å-1
PDB code 6MRSA
ΔdRMSD = -4.315Å
ΔGDT_TS = +0.07
F
α-helix
50
25
0.0
- 25
-10
0
ΔGDT_TS
ΔdRMSD
6O35A
low
high
helical content
PDB: 6O35
chain A
chain B
chain C
chain D
trRosetta prediction w/o energy minimization
ΔdRMSD = 3.64 Å
trRosetta
RGN2
bent helix
trRosetta prediction after energy minimization
ΔdRMSD = 0.33 Å
RGN2
trRosetta

**Supplementary Figure 4. (A)** Absolute performance metrics for RGN2 (purple) and trRosetta (salmon) across two sets of proteins with no known homologs. **(B)** Differences between RGN2 and trRosetta dRMSD values were computed for 35 *de novo* proteins. Points in left-top quadrant correspond to targets with negative ΔdRMSD and positive ΔGDT_TS, *i.e.,* where RGN2 outperforms trRosetta, and vice-versa for the right-bottom quadrant. The other two quadrants (white) indicate proteins where there was no clear winner. Only entries with <12Å ΔdRMSD are shown (all remaining entries lie in the quadrant where RGN2 outperforms trRosetta). **(C)** Stacked bar chart shows the relative fractions of different secondary structure elements in proteins where one method dominates. The total height of each bar indicates protein length. **(D)** (Top) trRosetta was able to better predict 6O35A, a non-covalently linked alpha-helical bundle monomer with a 0.111Å$^{-1}$ curvature. (Bottom) RGN2 consistently outperforms trRosetta on non-curved alpha-helical domain proteins interspersed by bends, turns, and beta strands (*e.g.,* 6MRSA and 6DG6A). **(E)** RGN2 was able to accurately predict the overall fold of *de novo* designed protein 6MRS (chain A) despite its high beta sheet content as the orientation and curvature of these beta pleats (B1-B4) are primarily controlled by straight alpha helices (H1 and H2) and hydrogen-bonded turns, both of which are strengths for RGN2. **(F)** Helical geometry of the monomer 6O35, whose structure is primarily dictated by interactions with neighboring monomers and interspersed stabilizing water molecules, as evidenced by the PDB structure. Raw trRosetta predictions capture these interactions and hence predict helical bends even for a disembodied monomer. However, Rosetta energy minimization of the trRosetta-predicted monomer removes such energetically unfavorable bends (in the monomer), recapitulating the geometry predicted by RGN2.

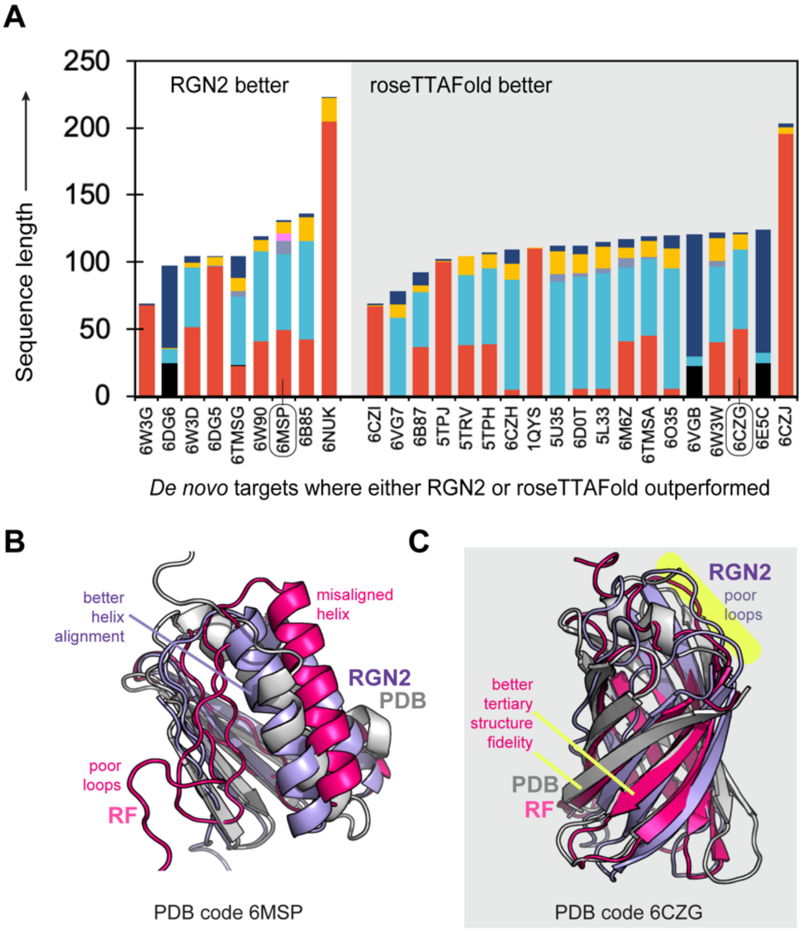


**Supplementary Figure 5. (A)** Stacked bar chart shows the relative fractions of different secondary structure elements in *de novo* proteins where RGN2 outperformed RF. The total height of each bar indicates protein length. **(B-C)** One case each where RGN2 outperforms RF and vice-versa.
