## Supplementary Text for "Single-sequence protein structure prediction using language models from deep learning"

***RGN2 performs better than trRosetta on Uniclust30 orphan dataset for longer proteins***

Alpha helices with tertiary structure elements most challenged RGN2. In **Supplementary Figures 2A-B** relative performance of RGN2 and trRosetta is compared based on secondary structure composition (**Supplementary Figures 2C**). Most structures with high helical content (HC) (both α-helices and $3_{10}$ helices), on which trRosetta outperformed RGN2 had high curvature $\kappa > 0.001Å^{-1}$, where $\kappa$ is the inverse of the radius of curvature of a helical domain). Four such proteins are shown in **Supplementary Figure 2D**, labelled by PDB accession code: 1D9J (HC – 47%, $\kappa \sim 0.11Å^{-1}$), 6CL3 (HC –100%, $\kappa \sim 0.09Å^{-1}$), 1WQD (HC –71.4%, $\kappa \sim 0.33Å^{-1}$), and 2KJF (HC – 73%, $\kappa \sim 0.118Å^{-1}$). It is unclear why curved helices are more challenging for RGN2, but it is likely that even though helical propensity is readily detectable from individual sequences subtler feature such as curvature is not. Similarly, all three MSA-based methods better predicted global fold (ΔGDT_TS > 24) of proteins with curved helices, RGN2 predictions were all within 1Å of them; in contrast when RGN2 outperformed these methods, it did so by a larger margin.

***RGN2 outperforms trRosetta on de novo dataset of 35 proteins***

In **Supplementary Figure 4A-C**, we illustrate RGN2's performance comparison on *de novo* designed proteins with trRosetta along with the structures of some notable predictions. In general, RGN2 accurately predicts helical packing and the geometry of connecting linkers as exemplified by structures 6MRSA and 6DG6A, but struggles with curved beta-pleats (*e.g.,* 6E5C) and the rare instances of bent helices (*e.g.,* 6O35A) (**Supplementary Figure 4D**). In contrast to the 6O35A target, the RGN2 prediction of monomeric 6DG6A conforms more closely to the experimental structure than that predicted by trRosetta (ΔdRMSD ~ 4.2Å) as the helices of 6DG6A show near-zero curvature ($\kappa \sim 10^{-3}$ $Å^{-1}$) and are linked by hydrogen-bonded turns, illustrating the high accuracy that RGN2 achieves with these elements. Structure 6MRSA further illustrates this—two straight alpha helices (H1 and H2—see **Supplementary Figure 4E**) span 48% of its residues while four beta strands (B1-4) cover most of the rest (37%). Despite the presence of a beta-pleat, its overall fold appears to be primarily stabilized by four electrostatic contacts between the helices and beta-strands, specifically the top and bottom of helices H1 and H2 and strands B1 and B4. Both of these helices are straight; RGN2 outperforms trRosetta on this protein with a ΔdRMSD of ~4.3Å.

One potential source of error in prediction arises from conformational changes that ae induced by multimerization: RGN2 is trained exclusively on monomeric proteins. In contrast, an MSA-based method such as trRosetta is expected to infer inter-domain contacts within homomultimers from the same co-variation signal used to detect intra-domain contacts. In 6O35A—a structure RGN2 predicts poorly— the geometry of several helices is strongly influence by tetramerization (stabilized by several water molecules). When we computed a monomeric conformation for 6O35A by using 1,000 steps of all-atom relaxation using Rosetta and explicit water molecules as solvent, we found that bends in the helices were lost (**Supplementary Figure 4F**), RGN2 and trRosetta predicted very similar structures (dRMSD ~1.7Å). We therefore conclude that RGN2 more closely resemble the monomeric state of 6O35A. Moving forward, encoding the presence of explicit water molecules, and other non-covalently bound molecules would enable detection of these bends in protein targets.
